# The *Seminavis robusta* genome provides insights into the evolutionary adaptations of benthic diatoms

**DOI:** 10.1101/2020.02.11.942037

**Authors:** Cristina Maria Osuna-Cruz, Gust Bilcke, Emmelien Vancaester, Sam De Decker, Nicole Poulsen, Petra Bulankova, Bram Verhelst, Sien Audoor, Darja Stojanovova, Aikaterini Pargana, Monia Russo, Frederike Stock, Emilio Cirri, Tore Brembu, Georg Pohnert, Per Winge, Atle M. Bones, Gwenael Piganeu, Maria Immacolata Ferrante, Thomas Mock, Lieven Sterck, Koen Sabbe, Lieven De Veylder, Wim Vyverman, Klaas Vandepoele

## Abstract

Benthic diatoms are the main primary producers in shallow freshwater and coastal environments, fulfilling important ecological functions such as nutrient cycling and sediment stabilization. However, little is known about their evolutionary adaptations to these highly structured but heterogeneous environments. Here, we report a reference genome for the marine biofilm-forming diatom *Seminavis robusta*, showing that gene family expansions are responsible for a quarter of all 36,254 protein-coding genes. Tandem duplications play a key role in extending the repertoire of specific gene functions, including light and oxygen sensing, which are probably central for its adaptation to benthic habitats. Genes differentially expressed during interactions with bacteria are strongly conserved in other benthic diatoms while many species-specific genes are strongly upregulated during sexual reproduction. Combined with re-sequencing data from 48 strains, our results offer new insights on the genetic diversity and gene functions in benthic diatoms.

## Introduction

In contrast to planktonic environments, benthic habitats are complex and heterogeneous, characterized by sharp and sometimes dynamic microscale gradients of light, oxygen, nutrient availability and redox state [1]. Benthic organisms are regularly exposed to inhospitable conditions such as extended periods of darkness, anoxia or the presence of toxic compounds such as sulfides [2]. Yet, illuminated surfaces in shallow aquatic systems are inhabited by dense and highly productive phototrophic biofilms which fuel food webs, modulate fluxes of carbon and nutrients, and can even stabilize sediments through the production of copious amounts of extracellular polymeric substances [3, 4].

In temperate and polar regions, phototrophic biofilms are frequently dominated by diatoms. These stramenopile microalgae are key players in aquatic ecosystems, accounting for up to 20% of global primary production [5]. They are uniquely characterized by a complex, bipartite silica cell wall and a particular size reduction-restitution life cycle [6]. Compared to the largely planktonic group of centric diatoms, the predominantly benthic and evolutionary younger pennate diatoms are far more species-rich [7]. This remarkable diversification is especially pronounced among the raphid pennates, being attributed to their heterothallic mating systems and active cell motility. Heterothally (differentiation in two or more mating types, as opposed to homothally in most centrics) promotes outcrossing, generating high levels of genetic diversity. Diatom cell motility, which is thought to be driven by an intracellular actomyosin cytoskeletal motor system linked to adhesive mucilage secretions from a cell wall slit termed the raphe, allows active positioning along chemical and physical microgradients within or on submerged surfaces. Importantly, motility of gametangial cells or gametes enables pheromone-guided mate finding, thus optimizing encounter success between opposite mating types in highly diverse and densely packed biofilms [8]. Combined, these two features may underlie the extraordinary diversity of raphid diatom species with often finely differentiated abiotic and biotic microniches [2].

However, to date, diatom genomic studies have mostly focused on planktonic centric species or on pennate species that colonize the planktonic environment. The first diatom genome to be sequenced was the planktonic centric *Thalassiosira pseudonana* [9], followed by the pennate *Phaeodactylum tricornutum* [10], which evolved morphological plasticity to switch between benthic and planktonic morphotypes. More recent studies have extended our knowledge of the complexity of diatom genomes, some examples being the oleaginous pennate *Fistulifera solaris* [11] with an allodiploid genome structure and the cold-adapted pennate *Fragilariopsis cylindrus* with a highly heterozygous genome showing allele-specific expression in response to environmental stresses [12]. While these first diatom genomes were small (27-61 Mb), sequencing of diatoms with larger genomes, including the centric *Thalassiosira oceanica* (92 Mb) [13] and the araphid pennate *Synedra acus* (98 Mb, also named *Fragilaria radians*) [14], have shown that these species have more than twice the number of protein-coding genes than the two first sequenced diatoms (*T. pseudonana* and *P. tricornutum*), suggesting the traditional diatom models are underestimating gene diversity.

In this work, we explore the genomic features of the pennate raphid benthic diatom *Seminavis robusta* to improve our understanding of genome evolution in diatoms as well as to provide insights into the molecular basis of ecological adaptations to the benthos. *S. robusta* resides in biofilms in shallow coastal habitats [15] and has been developed as a model system to study life cycle regulation, sexual reproduction and ecology of benthic pennate diatoms [16, 17]. Through the integrative analysis of the genome sequence of *S. robusta*, re-sequencing data of additional strains, detailed gene expression profiles, and gene information of 88 other diatoms, we shed light on key genes mediating differences in cell symmetry, motility and environmental adaptations between distinct groups of diatom clades.

## Results and Discussion

### Genome sequencing, assembly, gene annotation and expression atlas

Illumina paired-end sequencing in combination with long PacBio reads of the *Seminavis robusta* D6 strain, yielding 79× and 34× coverage, respectively, were used as input for genome assembly (*Supplementary Table S1*). A k-mer distribution analysis of the Illumina reads revealed a high level of heterozygosity (0.79%) and an estimated genome size of 117 Mb (*Supplementary Figure S1*), while flow cytometry yielded an estimate of 153 Mb. Both PacBio and Illumina data were used to generate several genome assemblies and to compare the performance of different tools and integration strategies, keeping as a final assembly the one that obtained the best balance between contiguity, completeness and quality (*Supplementary Table S2* and *Supplementary Note S1*). After removing four scaffolds corresponding to bacterial contamination (*Supplementary Figure S2*), the assembly consisted of 4,754 genomic scaffolds covering 125.57 Mb, which is within the range of estimated genome sizes, and had a scaffold N50 of 51 kb (*Figure 1A*). This genome assembly was further experimentally validated through the successful amplification of 22 regions that had the expected size (*Supplementary Figure S3*).

**Figure 1:**
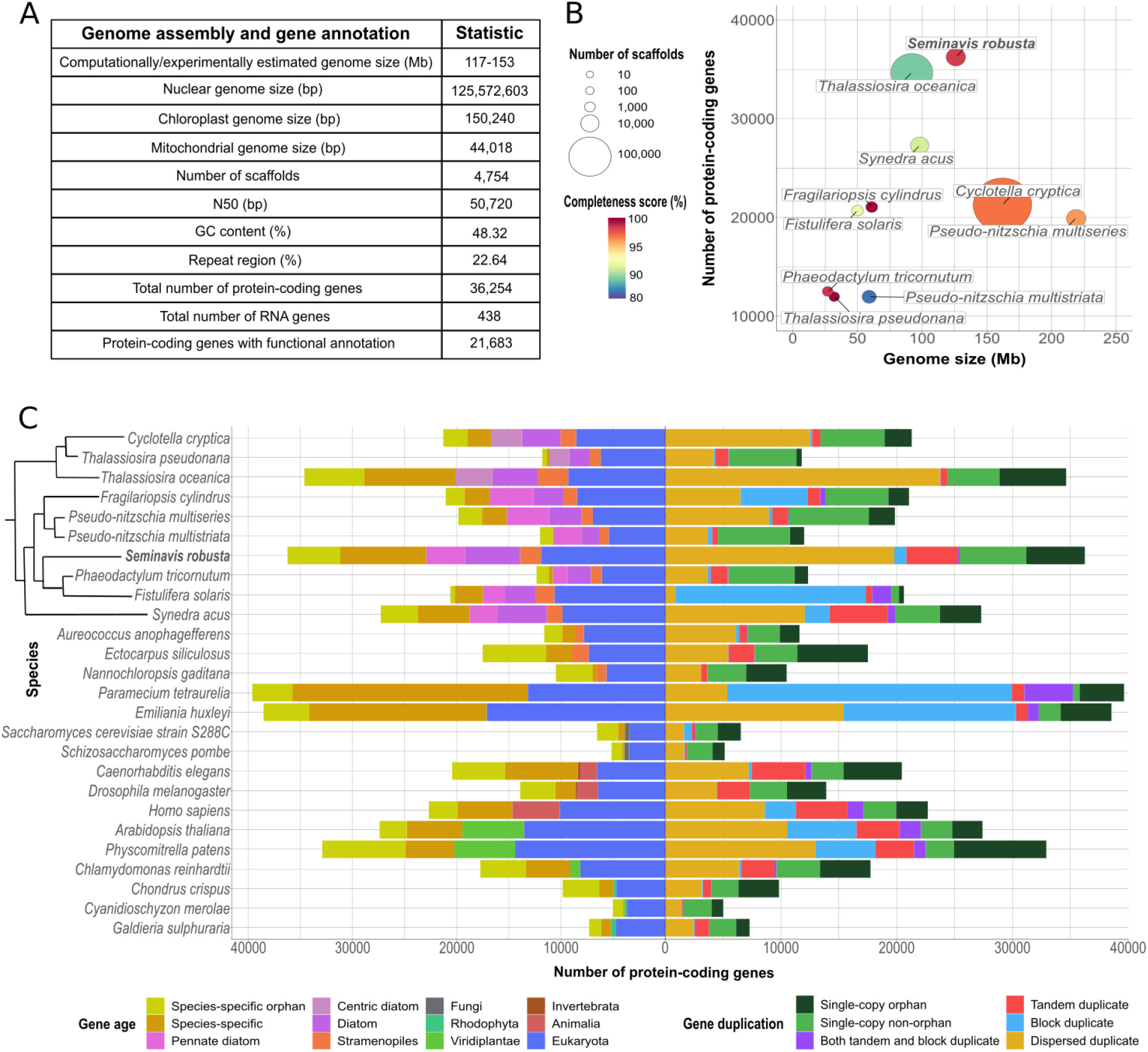
Genome properties for *S. robusta* and comparison with other sequenced diatoms. **(A)** Summary of the *S. robusta* genome assembly and gene annotation statistics. **(B)** Scatter plot showing genome assembly contiguity and gene family completeness score in sequenced diatom genomes. Every dot represents a diatom genome assembly, the x-axis displays the genome size in Mb whereas the y-axis represents the number of protein-coding genes. Genome assemblies are colored according to the gene family completeness score in a rainbow scale from blue to red. The size of the circle indicates the number of scaffolds in the genome assembly. **(C)** Comparative genomics analysis among diatoms and other eukaryote species. Left side of the bar plot represents the age of the genes inferred through phylostratification whereas the right side represents the duplication information. The phylogenetic relationship between diatom species is shown in a cladogram.

Repeat detection analysis revealed that 23% of the *S. robusta* genome assembly consists of repeats and transposable elements (*Supplementary Table S3*). After masking these regions, the *S. robusta* genome was subjected to an initial round of gene prediction and manual curation, followed by the use of newly generated expression data to improve gene models and identify additional expressed genes. Besides nuclear RNA gene prediction, resulting in 54 rRNAs, 227 tRNAs, 46 snoRNAs and 18 snRNAs, the chloroplast and mitochondrial genomes (*Supplementary Figure S4A-B*) were also annotated, leading to a total number of 36,254 protein-coding genes and 438 RNA genes (*Figure 1A*). To date, *S. robusta* is the diatom with the largest number of predicted protein-coding genes, of which 88% show expression support (*Supplementary Figure S5A*), 63% share similarity to proteins from other eukaryotes and 60% are functionally annotated. Application of a phylogeny-based horizontal gene transfer detection procedure identified 1,741 genes of putative bacterial origin [18]. Assessing the accuracy and quality of *S. robusta* gene models revealed the successful recovery of 99% of core gene families conserved in *Bacillariophyta* (*Supplementary Note S1*), demonstrating the higher completeness of the *S. robusta* gene annotation compared to other diatoms with large (> 90 Mb) genomes (*Figure 1B*).

Newly generated expression data combined with existing data were used to create a large-scale gene expression atlas profiling 31 different experimental conditions. This atlas comprises a total number of 167 samples (4,2 billion Illumina reads) and covers different sexual reproduction stages, abiotic stresses and bacterial interaction related treatments, the latter including responses to bacterial exudates as well as N-acyl homoserine lactones (AHLs), a class of signaling molecules involved in bacterial quorum sensing (*Supplementary Table S4 and Supplementary Figure S5B*). To investigate the functional significance of the *S. robusta* protein-coding genes, a differential expression analysis was performed, obtaining a total of 27,963 (77%) differentially expressed genes.

### Tandem duplication as a driver of gene family expansions for adaptive evolution

In order to study gene organization and evolution, we built the PLAZA Diatoms 1.0 platform (https://bioinformatics.psb.ugent.be/plaza/versions/plaza_diatoms_01/) to identify gene families based on the protein-coding genes from 26 eukaryotic genomes, which include nine other diatoms (*Supplementary Table S5*). Diatoms with the largest number of protein-coding genes (*S. robusta*, *T. oceanica* and *S. acus*, *Figure 1B*) have overall more families and a higher proportion of species-specific families compared to diatoms with smaller genomes such as *P. tricornutum* (*Figure 1C* and *Supplementary Note S2*). We identified 594 *S. robusta* expanded families (*Figure 2A*, *Supplementary Figure S6 and Supplementary Data S1*), driven by different duplication mechanisms (*Supplementary Figure S7A-B*), containing 9,178 genes. Despite the presence of a small number of block duplicates, no evidence for a recent whole-genome duplication was found. Several *S. robusta* expansions include families encoding for genes involved in molecular sensing, light signaling and motility (*Figure 2B* and *Supplementary Note S3*). For instance, we found the single-domain voltage-gated channels (EukCatAs) and the red/far-red light sensing phytochrome (DPH) families expanded by dispersed and tandem gene duplicates, respectively. While the EukCatAs family has been shown to modulate gliding locomotion in raphid pennate diatoms through fast Na^+^/Ca^2+^ signaling [19], the DPH family may be relevant for sensing the density and stress status of biofilms [20] (*Supplementary Note S3*).

**Figure 2:**
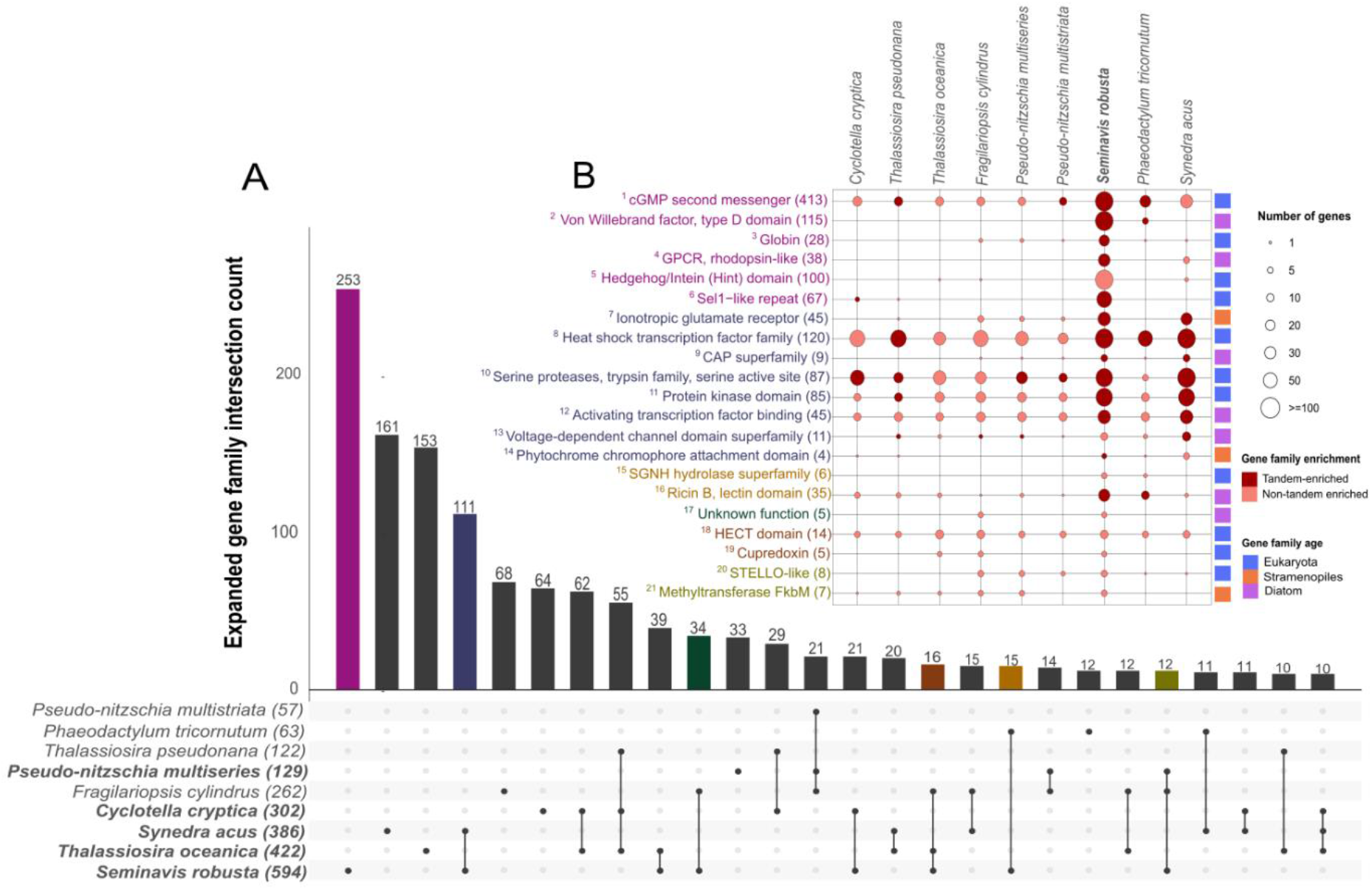
Species-specific and shared gene family expansions in diatoms. **(A)** Upset plot showing the intersection of gene family expansions in diatoms. Each row represents a diatom species with in parenthesis reporting the total number of expanded gene families. The barplot indicates the total gene family count in each intersection, displaying only intersections that contain at least ten gene families. Diatoms with a genome size > 90 Mb are highlighted in bold. **(B)** Examples of species-specific and shared gene family expansions in *S. robusta*. Each column represents a diatom species and each row a given gene family showing expansion in *S. robusta*, indicating the total number of genes in *S. robusta* in parenthesis and matching the font color with the intersection subset in panel A. The size of the circles is proportional to the number of genes falling under the given gene family per species whereas the color of the circles indicates if the gene family is significantly tandem-enriched. Numbers in superscript refer to families annotated in *Supplementary Data S1*.

*S. robusta* displays a remarkably high number of tandem duplicates (4,594 genes), which is also observed for *S. acus* (5,670 genes) as well as several multicellular eukaryotes (*Figure 1C*). A DNA coverage analysis indicated that the vast majority (81-84%) of these tandem gene duplicates, organized in gene cluster arrays, exist as distinct copies in the *S. robusta* genome and are not an artefact of the assembly procedure (*Supplementary Figure S8*). Tandem duplications have been shown to play an important role in accommodating responses to external stimuli and adaptation to rapidly changing environments [21] as well as generating novel expression patterns through exon shuffling [22]. Of the 318 expanded families in S. *robusta* having tandem duplicates, 69 were significantly enriched in tandem duplicates and only six of these families were shared with *S. acus* (*Figure 2B*), revealing that tandem-mediated family expansions are largely species-specific (*Supplementary Figure S9*). *S. robusta* tandem gene duplicates mainly consist of leucine-rich repeat (LRR) containing proteins, cyclases, heat shock factors, serine proteases, ubiquitin ligases, ricin B-like lectins, rhodopsins, ionotropic glutamate receptors, polyketide enzymes and globins, all genes that may be important for the adaptive evolution of this species (*Supplementary Table S6*).

As 70% of *S. robusta* genes are part of multi-copy gene families, we evaluated potential divergence and redundancy of gene duplicates by studying their expression profiles (*Supplementary Figure S10*, *Supplementary Data S1*). We observed that families with an increasing number of genes tended to display higher expression divergence than smaller families (*Figure 3A*), and that families showing expression divergence are significantly enriched in eukaryote and diatom age classes. Reversely, families showing expression conservation are significantly enriched in the species-specific age class (*Supplementary Figure S11A-B*). These results indicate that gene duplication increases expression divergence and that expression divergence increases with the age of gene duplicates. To determine under which conditions *S. robusta*’s families are expressed and in which processes they are involved, we identified families significantly enriched in upregulated genes under seven or more diverse experimental conditions (referred to as pleiotropic families) or in few specific related conditions (*Supplementary Data S1*).

**Figure 3:**
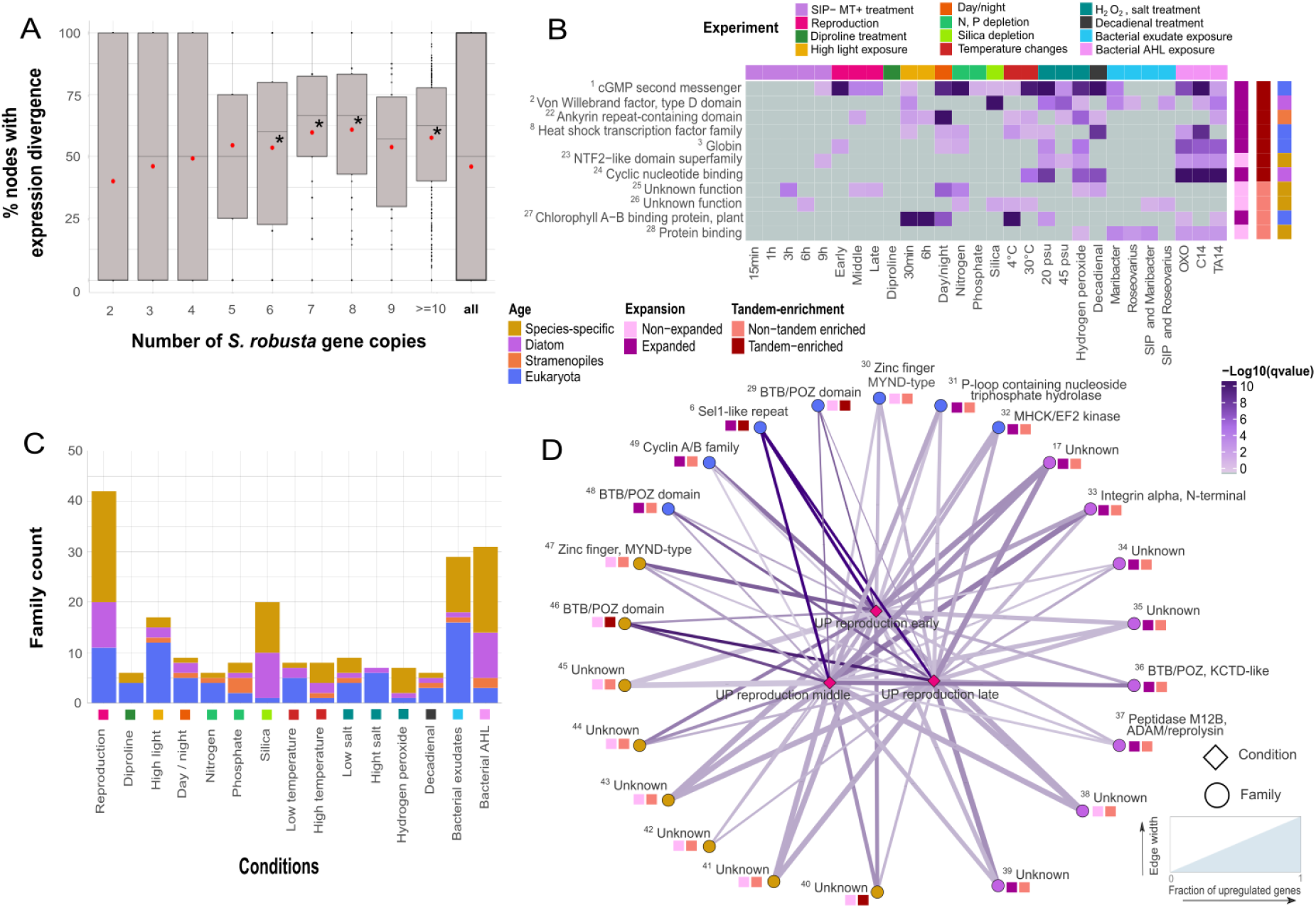
Expression analysis for *S. robusta* multi-copy gene families. **(A)** Expression divergence trend for multi-copy *S. robusta* families. The x-axis denotes the percentage of nodes showing expression divergence in the phylogenetic tree of the family, while the y-axis represents the number of *S. robusta* gene copies in the family. Average expression divergence percentages are indicated by red dots. Median expression divergence values significantly higher than the median of all nodes are highlighted with a star (pvalue<0.05, Wilcoxon rank sum test). **(B)** Heatmap showing pleiotropic families significantly enriched in upregulated genes for more than seven different conditions. The x-axis represents the different conditions/experiments whereas the y-axis reports the families. The significance of the upregulation in a certain condition for a family is shown in −log_10_(q-value) scale highlighted by a color gradient from grey to dark purple. Expansion, tandem enrichment and the age class of each family are highlighted in different colors on the right side of the heatmap. Numbers in superscript refer to families annotated in *Supplementary Data S1*. **(C)** Barplot showing family counts with significant condition-specific expression. The x-axis represents the different conditions/experiments whereas the y-axis represents the number of families having significant expression bias for that condition. The color of the bars denotes the family age distribution following the same color code as panel B. **(D)** Network showing families with significant specific expression in the three reproduction stages available. Families are represented with circles while conditions are represented with diamonds. The color of the circles denotes the family age and the color of the edges represents the significance of the upregulation, following both the same color-code as panel B. The edge width denotes the fraction of genes in the family that shows upregulation for the given condition. Expansion and tandem enrichment of each family are also highlighted following the same color-code as in panel B. Numbers in superscript refer to families annotated in *Supplementary Data S1*.

Eight out of the 11 pleiotropic families are expanded and/or enriched in tandem duplicates and several encode proteins involved in signaling (*Figure 3B*), indicating a strong link between gene family evolution, tandem duplication and *S. robusta*’s transcriptional response to environmental stimuli. The family that shows the highest pleiotropic signal is involved in cyclic guanosine monophosphate (cGMP) biosynthesis, which was shown to play a key role during the onset of sexual reproduction in *S. robusta* [16] as well as *Pseudo-nitzschia multistriata* [23]. A recent study suggests that this signaling pathway might also be involved in the response to bacteria [24]. Our results corroborate these functions, but also indicate that the cGMP-related signaling may have a much broader role than previously thought, showing significant upregulation in a wide-range of abiotic stresses (*Figure 3B*). For a heat shock transcription factor family (HSFs) expanded in diatoms [10] and showing pleiotropic expression in *S. robusta*, we identified this expansion happened through tandem duplication in several diatoms (*Figure 2B*). Interestingly, the *S. robusta* NTF2-like tandem-enriched pleiotropic family has 19/68 copies annotated with the polyketide cyclase SnoaL-like domain. Enzymes for polyketide metabolism are absent in some protist lineages but expanded in others such as dinoflagellates and haptophytes, suggesting that the evolution of these compounds may have played an important role in their ecological success [25].

von Willebrand factor, type D domains (vWDs) are found in numerous extracellular proteins and are believed to be involved in protein multimerization. For example, several adhesive proteins contain vWDs (e.g. zonadhesin, sea star foot protein, diatom adhesive trail proteins), that are thought to be important for maturation of the adhesive into multi-protein complexes [26, 27]. Sixty-one of the *S. robusta* vWD containing proteins also hold the diatom specific GDPH domain, hypothesized to be important for secreted diatom proteins with adhesive functions (e.g. motility, mucilage pads, gamete fusion) [27]. The identified pleiotropic tandem-enriched vWD family is highly abundant in raphid species and is significantly upregulated in 10 different conditions in *S. robusta*, half of these related to bacterial interactions, suggesting that they might be important for bacterial recognition, motility and adhesion. Hence, the expansion of the vWD family in *S. robusta* may reflect a key adaptation to highly heterogeneous benthic environments, which are inhabited by very diverse and dense bacterial populations compared to the water column.

In contrast to the wide expression of pleiotropic gene families, more than 200 families were identified showing significant enrichment for upregulation in one or a limited number of related conditions (*Supplementary Figure S12-13*). Notably, many of these families are species or diatom-specific and responsive to either sexual reproduction, silica depletion or bacterial interaction (*Figure 3C*, *Supplementary Note S4*). Hence, the strong specific expression of these families indicates a role in these distinctive diatom traits. Diatoms are unique by having a silica cell wall and a size reduction-restitution life cycle that for many diatoms also includes sexual reproduction [6]. Twenty-three out of the 42 reproduction responsive families are strongly expressed in the three profiled sexual stages (*Figure 3D*). The 12 families with functional annotation encode for proteins related to protein-protein interactions (BTB/POZ domain) [28], U box ubiquitin ligases (Sel1 repeats) and potential candidates for cell-cell recognition of gametangia and/or fusion of gametes (Integrins and M12B domain) [29]. In addition, the cyclin A/B family is also enriched in all stages of sexual reproduction, indicating that some of these genes play a specific regulatory role during meiosis (*Supplementary Figure S14*). Finally, eight of these reproduction responsive families show a simultaneous enrichment for hydrogen peroxide responsive genes (*Supplementary Figure S12A*) suggesting that reactive oxygen species signaling is important role during sexual reproduction.

*S. robusta* is found in shallow coastal habitats, often as part of subtidal biofilm communities, which can experience pronounced temperature changes. Seventy-one families showed specific expression towards bacterial interaction experiments (*Figure 3C* and *Supplementary Figure 12A-C*) and eight families towards high temperature (*Supplementary Figure S13*). Half of these families are species-specific and/or lack functional annotation, but their strong specific expression now sheds light on the biological processes they are putatively active in. The functionally annotated bacterial interaction responsive families are involved in intracellular signaling, oxygen sensing, detoxification, oxidative stress responses and cell adhesion. As an example, Arf GAPs can function as regulators of specialized membrane surfaces implicated in cell migration [30]. Together with their strong upregulation in the bacterial interaction experiments (*Supplementary Figure 12C*), we suggest that the tandem duplication driven expansion of these genes may be important for *S. robusta* cell adhesion and movement during biofilm formation. In contrast, the expansion by tandem duplicates of two families annotated as LRRs and ionotropic glutamate receptors (iGluRs) are relevant for *S. robusta*’s signaling during high temperature acclimatization (*Supplementary Figure S13*).

### Re-sequencing data reveals the conservation of gene family expansions and shows distinct patterns of gene selection

Re-sequencing data from 48 different *S. robusta* strains, representing three genetically distinct clades sampled from a small geographic region [15], was used to estimate the *S. robusta* pan-genome size and to validate the number of predicted protein-coding genes as well as the observed family expansions. While the core genome refers to genes present in all strains, the pan genome encompasses these core genes and dispensable ones present in only a subset of strains (*Figure 4A*). A total of 4,776 *de novo*-assembled genes were identified and after collapsing gene redundancy between different strains, 1,549 new dispensable genes absent from the reference genome were retained (*Supplementary Data S2-3*). Therefore, combined with the annotated genes in the reference genome, the *S. robusta* pan genome is estimated to cover 37,803 genes. Assessing the gene content diversity across all 49 *S. robusta* strains revealed that 74% of these genes corresponds to core genes. The remaining 26% represents the dispensable fraction, with 9,593 variable genes present in 2-48 strains and 90 specific genes present in only one strain (*Supplementary Figure S15*), the latter predominantly coming from the reference strain that had a higher sequencing depth (82/90, *Supplementary Data S4*). Globally, the re-sequencing of individual *S. robusta* strains indicates an average gene count of 35,012 protein-coding genes, therefore confirming the number of genes identified in the reference strain (*Figure 4B*). Inspecting our gene classification through short-read DNA gene length coverage analysis revealed that core genes had higher mean gene length coverage than dispensable genes (90% and 66%, respectively) (*Figure 4C*), while clustering strains by this gene coverage also recovered the three genetically distinct clades and hybrid strains previously described [15]. These results support our presence/absence variation classification and show this variability is not simply due to assembly or annotation errors.

**Figure 4:**
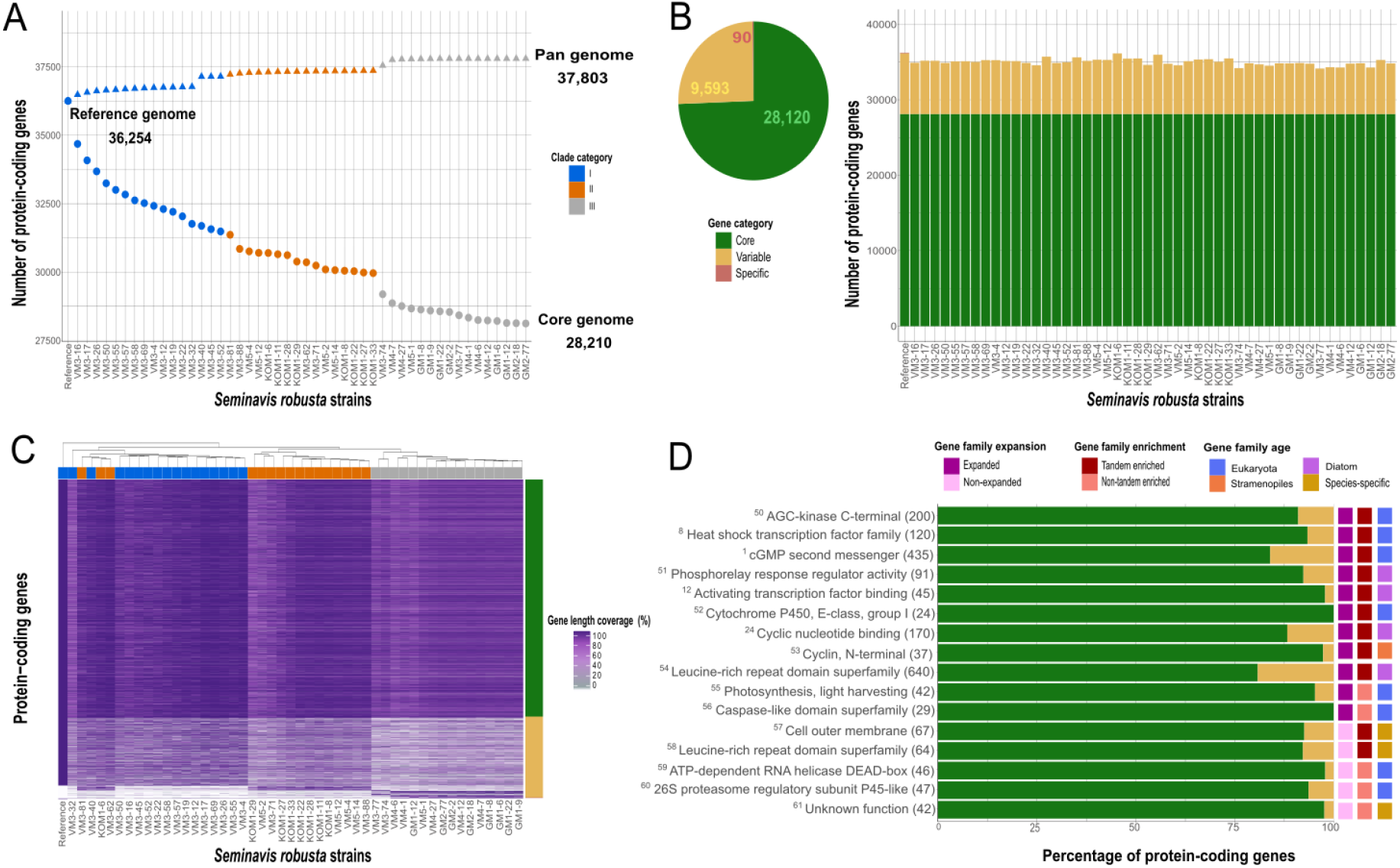
*S. robusta* within-species variability using a gene-based pan-genome analysis. **(A)** Representation of reference, core and pan gene size. The size of pan genome increases with each added strain up to 37,803 protein-coding genes whereas the size of core genome diminishes to 28,120 protein-coding genes. Clade category color-code refers to the population groups described in [15]. **(B)** Number of core, variable and specific genes per *S. robusta* strain. The pie chart shows the total gene count for each pan gene category (core, variable and specific). **(C)** Percentage of gene length coverage by short-read for all pan-genes for each strain. The x-axis represents the *S. robusta* strains whereas the y-axis represents all protein-coding pan genes. The percentage of horizontal gene coverage is highlighted by a color gradient from white (0%) to dark purple (100%). Pan-gene categories are labeled on the right side of the y-axis following the color code of panel B whereas clade categories are labeled on the upper part of x-axis following the color code of panel A. **(D)** Set of gene families that are significantly enriched in core genes. The x-axis represents the percentage of protein-coding pan genes that are core or dispensable, following the color code of panel B, while the Y-axis represents gene families, denoting in parenthesis the total number of pan genes belonging to that gene family (reference and *de novo* pan genes). Expansion, tandem enrichment and age of each family are highlighted in different colors on the right side of the y-axis.

Determining the size of the core gene content using re-sequencing data of ten geographically distant accessions in *P. tricornutum* revealed that 95% of the genes present are shared with all accessions (11,959/12,517 core genes, 172 *de novo*-assembled genes). The six times higher fraction of dispensable genes in *S. robusta* suggests much more *de novo* gene evolution in this benthic diatom compared to *P. tricornutum*, although convergent gene loss due to long-term culturing for the latter species might provide an alternative explanation. Nevertheless, this difference in genetic diversity is probably an underestimation given the large difference in geographic coverage of the strains analyzed for both species. The *S. robusta* dispensable gene content is however comparable with what has been described in the marine phytoplankton *E. huxleyi*, where 33% of the reference genes were missing in other strains [31], and plants (20-27% dispensable genes in *Brassica oleracea, Helianthus annuus*, *Brachypodium distachyon* and *Solanum lycopersicum* [32]). Future studies are needed to evaluate how variation in gene content differs between lab-cultured strains and isolates sampled from the environment.

In total, 73% of the reference tandem gene duplicates and 83% of the genes in expanded families are classified as core, indicating a strong conservation of duplicated genes in the different *S. robusta* strains. This finding was confirmed by the observation that 13 out of the 16 families significantly enriched in core genes were expanded and/or enriched in tandem duplicates (*Figure 4D*). These families are related to essential functions (e.g. light-harvesting involved in photosynthesis, caspases involved in proteolysis, cyclins controlling cell cycle regulation) but also encompass tandem-enriched expansions involved in signaling, including HSF, cGMP and LRR related proteins. Whereas many dispensable genes lack functional annotation, the remaining genes are enriched in DNA metabolism, zinc fingers, transposable element-derived proteins, and telomeric proteins (*Supplementary Data S4*). The latter are known to undergo rapid evolution, in particular, it has been shown that gene duplicates create telomere paralogs with novel functions in telomere replication and chromosome end-protection [33]. The observation that 63% of the *S. robusta* dispensable genes in the reference genome are expressed and several of these encode for transporters and metabolic proteins indicates this genomic variation represents transcriptionally active genes, potentially offering a template for phenotypic or physiological evolution in this diatom.

To evaluate conservation at the amino acid level and identify *S. robusta* genes that are under positive or purifying selection, we further used the re-sequencing strains to generate a high-confidence coding single nucleotide polymorphism (SNP) dataset (see *Supplementary Note S5*). The average estimates of nucleotide diversity at nonsynonymous (π_N_) and synonymous (π_S_) sites are 0.003 and 0.022 respectively. This indicates that two homologous sequences drawn at random from different strains will on average differ by 2.2% on synonymous sites and 0.3% on non-synonymous sites. The average π_N_ / π_S_ ratio is 0.14, which is smaller than what has been reported in other organisms such as *O. tauri* (0.20) [34], *C. reinhardtii* (0.22) [35], or the pennate diatom *P. tricornutum* (0.43) [36]. These differences reveal that globally the analyzed amino-acid composition of *S. robusta* genes undergoes a higher level of purifying selection, which is expected if it has a larger effective population size, as suggested by the higher π_S_. Furthermore, the distribution of the π_N_ / π_S_ ratio was estimated on individual genes (*Supplementary Data S4*). Interestingly, genes under positive selection are significantly enriched in genes upregulated during early, middle and late sexual reproduction stages, showing that genes involved in sexual reproduction are more prone to undergo diversification in their amino acid sequence, resulting in potential new protein functions. Although some of these genes belong to pleiotropic families (e.g cGMPs and vWDs) and families with reproduction expression bias (e.g. BTB/POZ domain and U box ubiquitin ligases) discussed previously, the vast majority are however genes of unknown function (100/128 genes) and/or species-specific single-copy (63/128), representing interesting candidates for future experimental characterization.

In addition, median π_N_ / π_S_ values were further analyzed to compare levels of purifying selection between distinct gene groups. Focusing on different gene age classes, we observed that gene age is positively correlated with purifying selection, with old genes being significantly more constrained than young genes (*Supplementary Figure S16A*). Genes with homologs in other eukaryotes are also more likely to be functionally annotated than species-specific genes, hence, poorly characterized genes show significantly less purifying selection than genes having functional annotations (*Supplementary Figure S16B*). Likewise, dispensable genes have significantly higher π_N_/π_S_ values than core genes (*Supplementary Figure S16C*), suggesting reduced functional constraint for variable genes in the *S. robusta* species complex. Strikingly, single-copy genes are not displaying significantly more purifying selection than duplicated genes. In contrast, when comparing between gene duplicate types, tandem duplicates are under less constraint than dispersed duplicates (*Supplementary Figure S16D*) indicating that tandem duplicates are more susceptible to amino acid changes, representing the potential evolution towards novel functionalities. We further observed that genes that are not expressed or show condition-specific expression are less constrained than genes that are broadly expressed (*Supplementary Figure S16E-F*), the latter which are frequently associated with housekeeping functions. Overall, our findings reveal that different levels of purifying selection act on different subsets of the *S. robusta* gene repertoire, and that dispensable genes as well as genes showing expression in few conditions are less evolutionary constrained, suggesting they might only be relevant in some specific environments.

### Signature genes for differences in cell symmetry, cell motility and environmental interactions between diatom clades

The Marine Microbial Eukaryote Transcriptome Sequencing Project (MMETSP) database [37] was used as a reference dataset to find *S. robusta* signature genes that might be implicated in specific morphological or ecological properties. These genes were identified by first determining, for each *S. robusta* gene, homologs in a set of 88 diatoms and subsequently selecting those genes predominantly present in pennate, raphid and benthic species, compared to centric, araphid and planktonic species, respectively (see *Material and methods and Supplementary Data S5-6*).

A significant number of genes showing high pennate signature are downregulated during silica depletion (55/630 genes). As many of these signature genes are related to cytoskeleton and membrane composition, this suggests they might play a role in cell wall and/or pennate cell shape formation (*Figure 5A*, *Supplementary Data S7* and *Supplementary Note S6*). Some examples of these cytoskeleton proteins include an actin-related protein 10 and a microtubule motor kinesin, modulating the dynein-mediated movement that contributes to for instance nuclear migration during mitosis [38]. Another gene with high pennate signature encodes a tubulin-tyrosine ligase/tubulin polyglutamylase, involved in the post-translational modification of tubulins which make up microtubules, as well as a CLASP N-terminal protein, implicated in the attachment of microtubules to the cell cortex and therefore regulating their stability. The loss of function mutation of the latter gene in *A. thaliana* has shown to result in various plant growth reductions, cell form defects and reduced mitotic activity [39], indicating that CLASP genes can potentially contribute to differences between pennate and centric diatoms in cell division and expansion. We also identified the nucleolar Las1-like protein showing high pennate conservation, which has been linked to cell morphogenesis and cell surface growth in yeast [40] and shows strong upregulation during early sexual reproduction in *S. robusta*. Related to membrane composition, we identified a CAAX amino terminal protease, inserted in the bilayer structure of the membrane, that is potentially implicated in protein and/or peptide modification and secretion [41]. The upregulation of this protease during early sexual reproduction hints towards a role in recognition during cell pairing between opposite mating types. In contrast, we found a fatty acid desaturase with high pennate signature upregulated during low temperatures, indicating that centric and pennate species might have also evolved different mechanisms to increase membrane fluidity for cold adaptation. The finding of 40 genes belonging to the CRAL-TRIO lipid binding superfamily further suggests the existence of differences in phospholipid metabolism [42].

**Figure 5:**
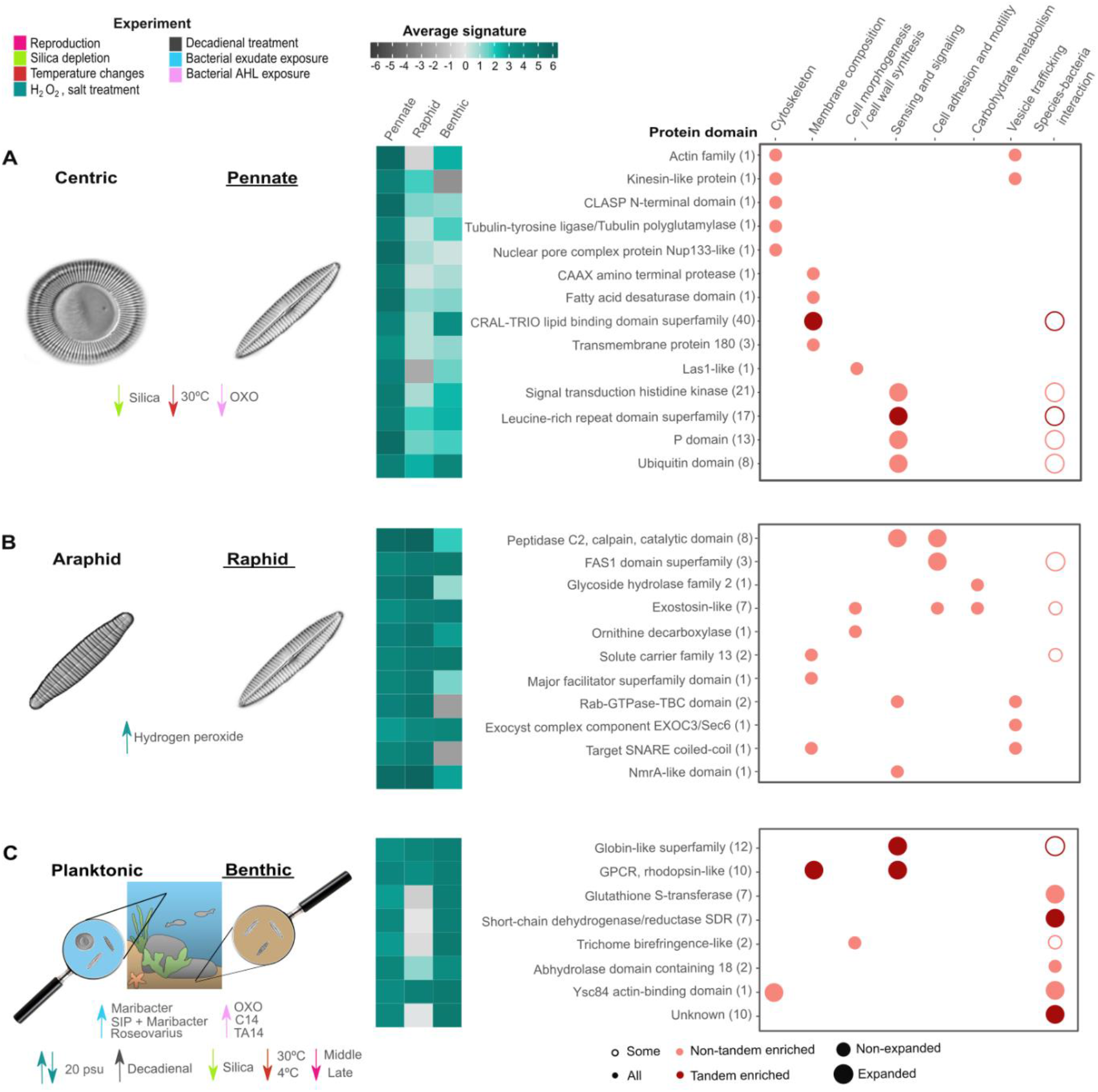
Selection of signature genes showing strong pennate (A), raphid (B) or benthic (C) conservation. Significant enrichment for differential expression in the genes showing the specified signature are highlighted with downward (for downregulation) or upward (for upregulation) arrows colored by experiment. Each row is a protein domain, the number of genes showing the signature and having that protein domain is indicated in parenthesis. If any of the genes containing one of the highlighted protein domains belong to an expanded and/or tandem enriched family, this is encoded by the size and color of the circles. The fill of the circles indicates if all or some genes with a given protein domain are upregulated during bacterial interaction experiments. The average pennate/raphid/benthic signature per protein domain is highlighted by a color gradient from dark grey (−6) to dark green (6). Images from a pennate raphid, pennate araphid and a centric diatom examples are courtesy of Spaulding, S., Edlund, M. (2008) in Diatoms of North America, available from https://diatoms.org/.

Within the 241 genes showing a high raphid signature, several genes related to cell adhesion and motility were found, including three proteins containing the ancient cell adhesion domain fasciclin (FAS1) and eight with a ‘peptidase C2, calpain’ domain (*Figure 5B*, *Supplementary Data S7* and *Supplementary Note S6*). Raphid species are responsive to intracellular calcium, playing a role in changing the speed and direction of locomotion [43]. Calpain proteins are calcium-responsive intracellular proteases that are implicated in the regulation of cell migration by controlling the dynamics of both integrin-mediated adhesion and actin-based membrane protrusion, enabling cell movement by modifying these adhesion sites [44]. To achieve their gliding motility and underwater adhesion, raphid diatoms secrete carbohydrate-rich extracellular polymeric substances (EPS) [27]. In line with this, multiple proteins with strong raphid conservation related to carbohydrate metabolism were also identified, comprising one glycoside hydrolase and seven exostosin-like proteins, the latter encoding glycosyltransferases involved in the synthesis of sulphated proteoglycans present in the extracellular matrix of mammalian cells [45]. Interestingly, sulphated polysaccharides/proteoglycans have been identified in the EPS and cell wall associated glycoproteins of raphid benthic diatoms such as *Stauroneis amphioxys* and *Craspedostauros australis* [46, 47]. The strong conservation of several of these exostosin-like proteins in different raphid benthic species (*Supplementary Figure S17*) hints towards their importance for EPS synthesis in these diatoms. Additionally, the presence of a raphid-specific ornithine decarboxylase, upregulated during reproduction, further suggests there are unique enzymes in raphid species controlling cell wall synthesis, since the inhibition of ornithine decarboxylase, which affects polyamine biosynthesis, has shown to result in a dramatic alteration of diatom silica structure [48]. Furthermore, several raphid-specific membrane transporters were found, as well as proteins containing the ‘Rab-GTPase-TBC’, ‘exocyst complex component EXOC3/Sec6’ or ‘target SNARE coiled-coil’ domain, the latter three encoding key components related to vesicle trafficking [49].

Benthic diatoms are frequently part of illuminated surfaces in shallow aquatic systems that are inhabited by dense and highly productive phototrophic biofilms which are characterized by highly variable oxygen and light conditions. To identify genes that can be linked with molecular components explaining the success of diatoms in these environments, we explored the *S. robusta* gene functions showing strong conservation in other benthic diatoms. Within the set of 492 benthic signature genes, we indeed observed a strong over-representation of different genes that can directly be related to oxygen and light signaling as well as diatom-bacteria interactions (*Figure 5C*, *Supplementary Data S7*). Twelve globin-like *S. robusta* genes with high benthic signature, organized in three different families, were identified. Globins have been found across all phyla of life, meaning that the oxygen transport function of vertebrate hemoglobins is a relatively recent adaptation and that the early globin functions were enzymatic and oxygen-sensing. Dissolved and particulate carbon and nutrient levels are higher within benthic habitats compared to the water column, increasing microbial numbers and heterotrophic activity. As a result, oxygen is quickly depleted, leading to a steep redox gradient in the sediment [50], which implies benthic species may have evolved mechanisms to sense oxygen concentration changes and regulate chemotaxis towards optimal conditions. One of the globin-like signature families is within the top pleiotropic families expanded by tandem duplication (*Figure 3B* and *Supplementary Figure S18*) and is restricted to pennate diatoms (*Figure 2B*), while the other two families encode for benthic-specific globins (*Supplementary Figure 19*). Whereas one of these benthic-specific families shows strong upregulation under hydrogen peroxide treatment, future experiments are needed to clarify the role of these globin-like proteins in sensing oxygen concentrations.

We identified ten putative G-protein-coupled receptor (GPCR) rhodopsin-like proteins having high benthic signature, of which seven belong to one family expanded by tandem duplication (*Figure 2B*). In the ocean, the spectral composition of light becomes progressively dominated by blue-green light (400-500 nm) with increasing depth. In biofilms, differential absorption of light by phototrophs and particulate and dissolved matter can contribute to variability in the spectral composition and intensity of available photosynthetically active radiation. Hence, benthic diatoms might have expanded and/or evolved their gene repertoire for light sensing. GPCR rhodopsin-like receptors transduce extracellular signals of a wide range of stimuli including light [51]. The existence of homologs to bacterial proteorhodopsins in some diatoms has been suggested to represent an alternative ATP-generating pathway, especially in iron-limited regions like the Southern Ocean [52]. A phylogenetic analysis revealed that the tandem-enriched family containing GPCR rhodopsin-like proteins with high benthic signature is not similar to these proteorhodopsins, rather, their members are more related to rhodopsin-like proteins found in other eukaryotes (*Supplementary Figure S20A*). Expression analysis of the GPCR rhodopsin-like protein ortholog in *P. tricornutum* shows this gene is specifically upregulated under blue-green light conditions (*Supplementary Figure S20B*), confirming our hypothesis this family plays an important role in light sensing.

The enrichment of benthic signature genes upregulated during bacterial interaction experiments (69 genes, *Supplementary Figure S21*) strongly correlates with a benthic lifestyle, where diatom-bacteria encounters are frequent due to high cell packing in biofilm communities. Secretion of EPS by benthic diatoms and their physicochemical properties are important factors driving biofilm formation, which has been shown to be strongly influenced by interaction with bacteria and/or their extracellular substances [53]. Here, we found one of the previously mentioned exostosin-like proteins, as well as a trichome birefringence-like protein, both potentially related to benthic EPS synthesis. In particular, the latter protein is responsible for the O-acetylation of polysaccharides in bacterial biofilm formation, suggesting O-acetylation may also play an important role in benthic diatom biofilm architecture [54]. Benthic signature genes contain several NAD- or NADP-dependent oxidoreductases and detoxification enzymes (*Supplementary Figure S21)*, covering seven short-chain dehydrogenases/reductases (SDRs) and seven glutathione S-transferases (GSTs). While SDRs are enzymes of a great functional diversity, GSTs are often involved in the detoxification of reactive oxygen species (ROS) and xenobiotics, indicating that benthic diatoms have an evolved gene set to control ROS homeostasis and increase oxidative stress defense in biofilms [55].

### Concluding remarks

The reported *S. robusta* genome sequence and its integration with re-sequencing data from 48 additional strains, large-scale expression profiling, and transcriptome data from different classes of diatoms, provided a unique opportunity to shed light on the genome complexity and evolution in benthic diatoms. To date, *S. robusta* is the diatom with the largest number of protein-coding genes, of which 40% are of unknown function and 60% are diatom or species-specific. The strong specific expression pattern of many of these genes has provided valuable new insights into the biological processes they are involved in, including 42 families related to sexual reproduction and 71 families showing specific responses to bacterial interactions. Our findings therefore reveal new candidates controlling specific diatom traits. We show that the large number of *S. robusta* genes that originated through gene family expansions are conserved in the *S. robusta* species complex and are extending the gene repertoire of this species for molecular sensing, light signaling and motility. Tandem gene duplication is a prominent feature of pleiotropic gene families showing a complex pattern of wide-spread regulation in numerous conditions. Adaptation to the benthos through tandem gene duplication has also been observed in the penaeid shrimp [56], which shares with *S. robusta* the iGluR expansion. In particular, rhodopsin-like proteins, globins and detoxification enzymes seem to be key players in the ecological adaptation of not only *S. robusta*, but also of other benthic diatoms that dominate marine biofilms. Whereas the large fraction of dispensable genes in *S. robusta* indicates a major difference in gene space variation compared to *P. tricornutum*, the phenotypic, ecological and physiological consequences of missing genes in specific strains remain to be seen. Future studies of additional diatom species will help to identify and better define their adaptations to the benthic lifestyle, paving the road to understand the extraordinary evolutionary success of the pennate diatoms.

## Materials and methods

### DNA extraction, library preparation and genome size estimation

Both Illumina and PacBio technologies were used for sequencing of *S. robusta* D6 strain mating type plus (MT+) to take advantage of short-read (better quality) and long-read (better contiguity) sequencing approaches. In the case of Illumina, DNA was extracted using a DNeasy Plant Mini Kit (Qiagen), paired-end libraries were prepared with an insert size of 500-800 bp and sequencing was performed on a 2× 300bp Miseq system. In the case of PacBio, DNA was extracted with the CTAB method, libraries were prepared with an insert size of 10,000 bp, and sequencing was performed in two different locations: 10 SMRT cells were sequenced on the RSII at VIB Nucleomics Core (Leuven, Belgium) whereas an extra 1 SMRT cell was sequenced on the RS at GATC Biotech AG (Konstanz, Germany). All statistics related to DNA sequencing data and preprocessing are summarized in *Supplementary Table S1*. The genome size of *S. robusta* was experimentally estimated to be 153 Mb by flow cytometry. Additionally, k-mer distribution statistical approach was used to estimate the genome size (117 Mb), repeat content (23 Mb), and heterozygosity (0.78%) (*Supplementary Figure S1*).

### Genome assembly and repeat identification

Due to the complexity caused by the heterozygosity, several genome assemblies were generated to compare the results of different tools and assembly integration strategies, including Illumina-only, PacBio-only as well as hybrid approaches combining the Illumina and PacBio data (*Supplementary Table S2* and *Supplementary Note S1*). The ultimate chosen genome assembly was computed using the following protocol: i) Illumina paired-end reads were assembled into contigs using the haplotype-aware Platanus v1.2.4 [57] tool ii) a heterogeneity cleaning process was executed to identify potential redundant sequences still present in the assembly based on a pairwise comparison using BLASTn v2.3.0 [58] (>85% identity, >75% coverage), keeping the largest sequence as non-redundant (strategy similar to redundans pipeline [59]) iii) an extra round of scaffolding and gap-closing steps with Platanus was applied iv) PacBio reads were integrated using PBJelly v15.8.24 [60] v) a second heterogeneity cleaning process was performed onto the generated hybrid assembly vi) a final re-scaffolding with PBJelly was executed. The redundant sequences resulting from the heterogeneity cleaning comprised a total of 12.07 Mb. One third of the Illumina paired-end reads that did not map to the reference genome did map to these sequences, revealing that 94% of all Illumina reads are present in the assembled sequences.

Both *de novo*-based and homology-based approaches to mask repeats prior to gene prediction were employed by using RepeatModeler and RepeatMasker [61]. A summary of the detected repeats in the *S. robusta* genome and the detailed protocol can be found in *Supplementary Table S3*.

### Gene prediction and functional annotation

Reads from ten different RNA-Seq experiments in *Seminavis robusta* that were available when this analysis started (see *Supplementary Table S4*), were employed as a training to infer gene predictions using the BRAKER1 pipeline [62]. The resulting gene prediction contained 35,265 protein-coding genes and was named as gene annotation v1.0. To further improve the gene annotation, manual curation of the *S. robusta* expanded gene families (582 curated genes in total) was performed using the ORCAE [63] interface (gene annotation v1.1). Next, reads from 21 newly generated RNA-Seq samples were trimmed (see *Supplementary Table S4*), and mapped to the genome assembly using STAR v 2.5.2 [64]. These alignments were used to run the Genome-guided Trinity *De novo* Transcriptome v2.6.6 and PASA v2.3.3 pipeline [65] in order to add untranslated regions (UTRs) (to 14,634 genes) and new detected expressed gene models (938 genes), generating the final gene annotation v1.2 with a total number of 36,254 genes. The assessment of the quality of these gene models is described in the *Supplementary Note S1*.

Functional annotation of the protein-coding genes was performed by integrating the results from three different approaches. First, InterProScan v5.31 [66] was run to scan our sequences for matches against the InterPro protein signature database. Second, eggNOG-mapper v1 [67] was executed with DIAMOND mapping mode, based on eggNOG 4.5 orthology data [68]. Third, a consensus functional annotation per gene was computed based on similarity searches following the AnnoMine pipeline described in [69].

Non-coding RNA genes were predicted using Infernal v1.1.2 [70] (e-value<10e−03, coverage >=90%). The predictions that overcame the defined thresholds were manually checked in the Rfam database for the taxonomic origin, retaining mainly eukaryotic hits. In the specific case of tRNA genes, tRNAscan-SE v 1.31 [71] was also executed and the results were merged with the Infernal output.

### *S. robusta* comprehensive expression atlas and gene differential expression analysis

Cultures of *S. robusta* were subjected to 31 different conditions (167 samples) encompassing sexual reproduction, abiotic and biotic stress. New RNA-Seq data was generated for 83 samples and was complemented with previously published datasets. The novel transcriptomic data was produced by subjecting *S. robusta* cultures to various treatments followed by RNA extraction and Illumina sequencing. A detailed overview of conditions and procedures can be found in *Supplementary Table S4*.

The sequenced reads were mapped to the longest isoform of *S. robusta* gene models with UTRs using Salmon v0.9.1 [72]. Gene-level abundances of all 167 samples were imported in R using the tximport package v1.8.0 and Transcript Per Million (TPM) values were calculated. A gene was considered to be expressed when TPM >= 2 and to be condition-specifically expressed when Tau >= 0.9 [73]. For the differential expression analysis, genes with very low counts were filtered, retaining only genes with > 1 count per million (CPM) in at least three samples. Out of a total of 36,254 genes, 32,273 genes were retained after filtering. Counts for each experiment were loaded into a separate DGEList object from the R package EdgeR [74]. TMM normalization factors were calculated that are used as an offset to correct for differences in sequencing depth and RNA composition. After estimating tagwise dispersion, negative binomial GLMs were computed for every gene and contrast matrices were designed to define 31 conditions (see *Supplementary Table S4*). To test for differential expression, likelihood ratio tests (LRT) were carried out. To limit the number of differentially expressed genes, we tested for differential expression relative to a log_2_(fold change) threshold of ±1 with the glmTreat() function in EdgeR. FDR adjusted p-values were calculated for each comparison using the Benjamini-Hochberg correction, setting a p-value < 0.05 threshold for differential expression.

### Comparative genomics analysis

All *S. robusta* predicted genes were loaded into a diatom-centered version of the PLAZA comparative genomics platform [69], containing 26 eukaryotic genomes (*Supplementary Table S5*). A detailed list of tools used to build PLAZA Diatoms 1.0 is found in *Supplementary Table S7*. First, an all-against-all protein sequence similarity search was performed to subsequently delineate gene families. Families with more than three members and less than one thousand (from all species) where subjected to a multiple sequence alignment, followed by filtering and trimming and a phylogenetic tree construction using an approximate maximum likelihood method. Nearly single-copy families, having at least one and at most two gene copies per species, recovered across all diatoms were selected and the filtered and trimmed multiple sequence alignments for these families were concatenated per species (913 families, 269,486 amino acid positions). This alignment was used to infer the diatom species tree. Expanded families were delineated by calculating the Z-score profile of the gene copy number across all diatoms excluding the allodiploid *F. solaris*. Families where the variance is larger than two and the Z-score for a particular species is larger than three, were deemed expanded in that species. To assign genes or families to age classes, the taxonomic scope of each gene/family was evaluated using phylostratification [75]. Gene families significantly enriched in upregulated genes for a specific condition were computed using hypergeometric distribution with a q-value cutoff of 0.05 and a minimum of two hits. The same protocol was applied for computing families significantly enriched in core/dispensable genes (see next section). Gene duplication type and collinear regions, i.e. regions with conserved gene content and order, were also identified. Block duplication, also known as segmental duplication is the duplication of a whole genomic fragment. Here, a block is defined as consisting out of at least 3 anchor pairs, where a pair are genes belonging to the same gene family, which are separated by maximal 15 genes (gap size) and blocks are separated by at least 15 genes (cluster gap) when belonging to the same scaffold. Tandem genes on the other hand belong to the same gene family and form gene cluster arrays, that are located within 15 genes of each other (tandem gap). Dispersed duplicates are duplicated genes within a gene family, but are not found to have originated through either a block or a tandem duplication event. Genes categorized as tandem duplicates were validated through a DNA coverage analysis (*Supplementary Figure S8A-B*) and InterPro domain significant enrichment was calculated using the same protocol described before for differentially expressed genes (*Supplementary Table S6*).

### Identification of core, variable and specific *S. robusta* protein-coding genes

Adaptor-trimmed paired-end 151 bp reads were obtained from the Nucleomics Core Facility (VIB, Leuven, Belgium) for 48 different *S. robusta* strains. Quality of these reads was assessed using FastQC v0.11.4 and then they were uniquely aligned to the D6 reference genome using the BWA-MEM v0.7.5a tool [76], including nuclear, chloroplast and mitochondrial scaffolds. Next, reference gene presence/absence variation was characterized using the SGSGeneLoss package [77] (*Supplementary Data S4*), defining as gene absence when the horizontal coverage across all exons of the gene was < 5%.

*De novo* assembly of unmapped reads was conducted using Velvet as implemented by VelvetOptimizer [78] with a range of k-mers between 21 and 51 to enable the assembly of contigs from low coverage data. All contigs were aligned to the NCBI protein database and the diatom proteins from the PLAZA Diatoms 1.0 and MMETSP [37] using DIAMOND in translated DNA mode [79] (e-value < 10e−05, bitscore >= 200, --more-sensitive) to search for regions that potentially contain *de novo*-assembled dispensable genes. In addition, contigs were aligned to the D6 reference genome using BLASTn v2.7.1 [58] (>=75% identity and e-value < 10e−03) to identify regions that after *de novo* assembly were already represented in the reference genome. These regions were queried using BEDTools v2.27 [80] to subtract *de novo*-assembled genes that did not overlap with any of the identified reference regions. The resulting genes were screened for contamination by checking the taxonomy of the top 5 best hits in the DIAMOND search against the NCBI protein database, discarding genes exclusively matching non-eukaryotic genes. The remaining *de novo*-assembled genes were functionally annotated by transferring the functional annotation of its best hit in the PLAZA Diatoms 1.0 (*Supplementary Data S3*) and genes annotated as transposable elements were discarded. All these steps were executed for each *S. robusta* strain separately and a summary of these results can be found in *Supplementary Data 2*. To identify redundant *de novo*-assembled genes, the gene DNA sequences were processed using CD-HIT [81], with a similarity threshold of 95%, keeping the longest sequence of each cluster and generating a final non-reference dispensable gene dataset. The unmapped reads were mapped back to the *de novo*-assembled contigs that contained this final non-reference gene dataset in order to determine the presence/absence variation across all re-sequencing strains following the previously reported SGSGeneLoss protocol. All these mentioned steps were also applied to determine the presence/absence of genes in the *P. tricornutum* species complex as well as to identify extra *de novo*-assembled dispensable genes using resequencing data of 10 geographically distant *P. tricornutum* strains [36].

### SNP calling and analysis of gene selection patterns

Picard tool v1.94 was employed to mark duplicate reads in the previously-mentioned alignments of the 48 different *S. robusta* strains to the reference D6 genome. SNP calling was executed using Genome Analysis Toolkit (GATK) v3.7-0 [82] according to the GATK best practices. Variants were called per sample using HaplotypeCaller in GVCF mode and then passed together to the GenotypeGVCFs tool to generate a joint-called set of SNPs. The SNPs were further filtered out to generate a high confidence coding SNP dataset as explained in *Supplementary Note S5.* This high confidence coding SNP dataset comprised 28 Mb covering 20,891 genes and was functionally annotated using snpEff v4.3t [83]. Finally, we calculated the nucleotide diversity at nonsynonymous (π_N_) and synonymous (π_S_) sites and the ratio of these two for this set of genes by computing the polymorphism of the callable coding sequence of each gene, combining this with the functional information of snpEff output and correcting for the allele frequency of the mutation in the population (*Supplementary Data S3*).

### Identification of S. robusta genes with high pennate/raphid/benthic signature

All *S. robusta* protein-coding genes (core and dispensable) were aligned to the diatom MMETSP [37] proteins using DIAMOND in translated DNA mode [79] (--evalue 1e−05 --min-score 200 -- max-target-seqs 2000 --more-sensitive). These 180 diatom MMETSP samples were classified as pennate/centric, raphid/araphid and benthic/planktonic species, recovering protein sequences for 32 pennate, 56 centric, 18 raphid, 14 araphid, 8 benthic and 73 planktonic species in total (*Supplementary Data S5*). For benthic/planktonic classification, we only kept those species with a clearly benthic or planktonic lifestyle, excluding those that were unclear as well as the ice benthic diatoms. For each *S. robusta* gene, we computed a pennate, centric, raphid, araphid, benthic and planktonic score based on the number of species from each category that the *S. robusta* query gene had a homologous hit. This score goes from zero to one, where, for example, a pennate score of zero reports the gene has no hits with any pennate species, and a pennate score of one means homologs were found in all pennate species of the MMETSP samples. Next, we calculated the pennate/raphid/benthic signature as follows:

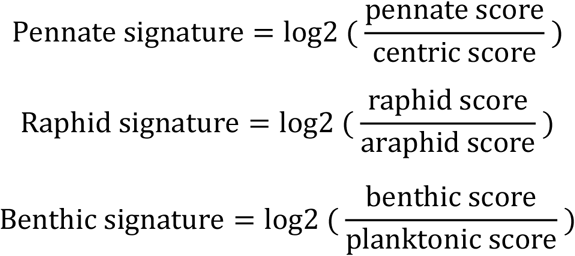

Since we are working with transcriptomes, note we cannot irrefutably affirm the absence of a certain gene in a diatom clade. As a consequence, to define genes with high pennate or raphid signature we used a signature threshold >=3 but also verified if the genomic data supports the observed signal, i.e. the diatom homologs of the family of the query gene were all from pennate and/or raphid diatoms. In contrast, as we only have genomic data from one true benthic diatom species (*S. robusta*), genes with high benthic signature were defined as having a signature threshold >=3 and a minimum of two benthic species in the MMETSP samples (*Supplementary Data S6*). Enrichment analysis of genes with a specific high signature were performed using the previously described enrichment analysis (q-value cutoff of 0.05 and minimum of two hits).

To evaluate how gene signature values change when using different cutoffs to identify homologs in the MMETSP dataset, we performed a control experiment using a more relaxed threshold (--evalue 1e−03 --min-score 0). This analysis revealed that 81%, 73% and 97% of the genes with high pennate, raphid and benthic signature had the same positive signature trend using both the stringent and relaxed cutoff, and that the use of these more relaxed thresholds leads to the recovery of more distantly related homologs that do not strongly change the detected signature (*Supplementary Figure S22A-C*).

## Data availability

Genome sequences data that support the findings of this study have been deposited in the European Nucleotide Archive with the accession code of PRJEB36614 while the fastq files representing the raw RNA sequencing data have been deposited in EBI Array Express under accession number E-MTAB-8685 (#currently password protected#). The genome sequences can also be downloaded from ORCAE platform (https://bioinformatics.psb.ugent.be/orcae/overview/Semro). Gene families are accessible in the PLAZA Diatoms 1.0 (https://bioinformatics.psb.ugent.be/plaza/versions/plaza_diatoms_01/), containing on each gene family page a direct link to the TPM and the differential gene expression matrix of all *S. robusta* genes. Results comprising expansions, tandem enrichment and expression divergence/conservation of *S. robusta* gene families as well as gene presence/absence variation, π_N_ / π_S_ values and signature genes are available in *Supplementary Data* S1–7.

## Acknowledgements

We would like to thank Sara Moeys and Jeroen Gillard for technical support, Luke Noble for his help with the Platanus assembly, Elisabeth Veeckman and François Bucchini for their help for assessing the gene completeness and Michiel Van Bel for his assistance with the PLAZA platform. CM.O-C., E.V., L. DV. and K. V. want to acknowledge the funding obtained by the BOF project GOA01G01715, G.B. to the Research Foundation Flanders (FWO) Aspirant grant (3F001916), N. P. to the Deutsche Forschungsgemeinschaft (PO 2256/1-1), P. B. to the Erwin Schrödinger fellowship from Austrian Science Fund (FWF) (J3692-B22), E.C., G. P. and F. S. to the European Union’s Horizon 2020 research and innovation program under the Marie Sklodowska-Curie grant agreement No. 642575 and MI. F. to the Gordon and Betty Moore Foundation through grant #7978.

## Contributions

K. V., W. V. and L. DV. Initiated and managed the *Seminavis robusta* genome sequencing project. P.B. extracted and prepared DNA sequencing and CM. O-C. performed genome assembly, gene predictions and functional annotation with the help of B. V. Manual gene curation was performed by CM. O-C., G. B., S. DD., P.B., S. A., D. S., A. P., L. DV. and K. V., aided by L. S. PLAZA Diatoms 1.0 was built by E.V. and gene family expansions were delineated together with CM. O-C. RNA-Seq experiments were designed and executed by G. B. and CM. O-C., while G. B. performed differential gene expression analysis. CM. O-C. studied gene family expression evolution. S. DD. generated the re-sequencing data and CM. O-C. processed these to determine gene presence/absence variation in *S. robusta* species complex. S. DD., CM. O-C., E.V. and G. P. were involved in the π_N_ / π_S_ analysis. CM. O-C. integrated MMETSP dataset with the help of K. S. and W. V., and calculated signature genes. MI. F., M. R., N. P., E. C., G. P., F. S., A. B., P. W. and T. M. analyzed specific gene families. CM. O-C. and K. V. wrote the manuscript with the help of all collaborators. All authors read and approved the final manuscript.

## Ethics declarations

